# Neuronal population mechanisms of lightness perception

**DOI:** 10.1101/294280

**Authors:** Douglas A. Ruff, David H. Brainard, Marlene R. Cohen

## Abstract

The way that humans and animals perceive the lightness of an object depends on its physical luminance as well as its surrounding context. While neuronal responses throughout the visual pathway are modulated by context, the relationship between neuronal responses and lightness perception is poorly understood. We searched for a neuronal mechanism of lightness by recording responses of neuronal populations in monkey primary visual cortex (V1) and area V4 to stimuli that produce a lightness illusion in humans, in which the lightness of a disk depends on the context in which it is embedded. We found that the way individual units encode the luminance (or equivalently for our stimuli, contrast) of the disk and its context is extremely heterogeneous. This motivated us to ask whether the population representation in either V1 or V4 satisfies three criteria: 1) disk luminance is represented with high fidelity, 2) the context surrounding the disk is also represented, and 3) the representations of disk luminance and context interact to create a representation of lightness that depends on these factors in a manner consistent with human psychophysical judgments of disk lightness. We found that populations of units in both V1 and V4 fulfill the first two criteria, but that we cannot conclude that the two types of information in either area interact in a manner that clearly predicts human psychophysical measurements: the interpretation of our population measurements depends on how subsequent areas read out lightness from the population responses.

**New & Noteworthy:** A core question in visual neuroscience is how the brain extracts stable representations of object properties from the retinal image. We searched for a neuronal mechanism of lightness perception by determining whether the responses of neuronal populations in primary visual cortex and area V4 could account for a lightness illusion measured using human psychophysics. Our results suggest that comparing psychophysics with population recordings will yield insight into neuronal mechanisms underlying a variety of perceptual phenomena.

## Introduction

A fundamental building block of our perception of object properties is *lightness*. For achromatic objects, lightness is the perceptual attribute of the object’s surface that varies from black, through gray, to white. The perceived lightness of an object’s surface is related to, but not completely determined by, its *luminance*, which characterizes the light reflected from the object to the eye. The luminance of the reflected light is the total spectral radiance after weighting by a measure of the spectral sensitivity of the visual system.

When only the diffuse surface reflectance of an object is varied, while its shape, position, the objects around it, and the incident illumination are held fixed, variation in luminance predicts variation in perceived lightness. When the context within which the object is varied, however, luminance is no longer a reliable predictor of lightness. Effects of context upon lightness support *lightness constancy*, wherein the perceived lightness of an object remains roughly constant across changes in illumination (Adelson, 2000; Kingdom, 2011).

The dissociation between physical luminance and perceived lightness is readily illustrated by Adelson’s checker shadow illusion, as elaborated by Gilchrist (http://persci.mit.edu/gallery/checkershadow; Gilchrist, 2006). Figure 1 illustrates our variant of this illusion. Disks that have the same luminance and the same immediate surround (and thus the same local contrast) have different perceived lightnesses; disks that are perceived as lying in a shadow (right panel of Figure 1, which we refer to as ‘shadow’ stimuli) are perceived as lighter than disks that do not lie in a shadow (left panel of Figure 1, which we refer to as ‘paint’ stimuli).

**Figure 1.**
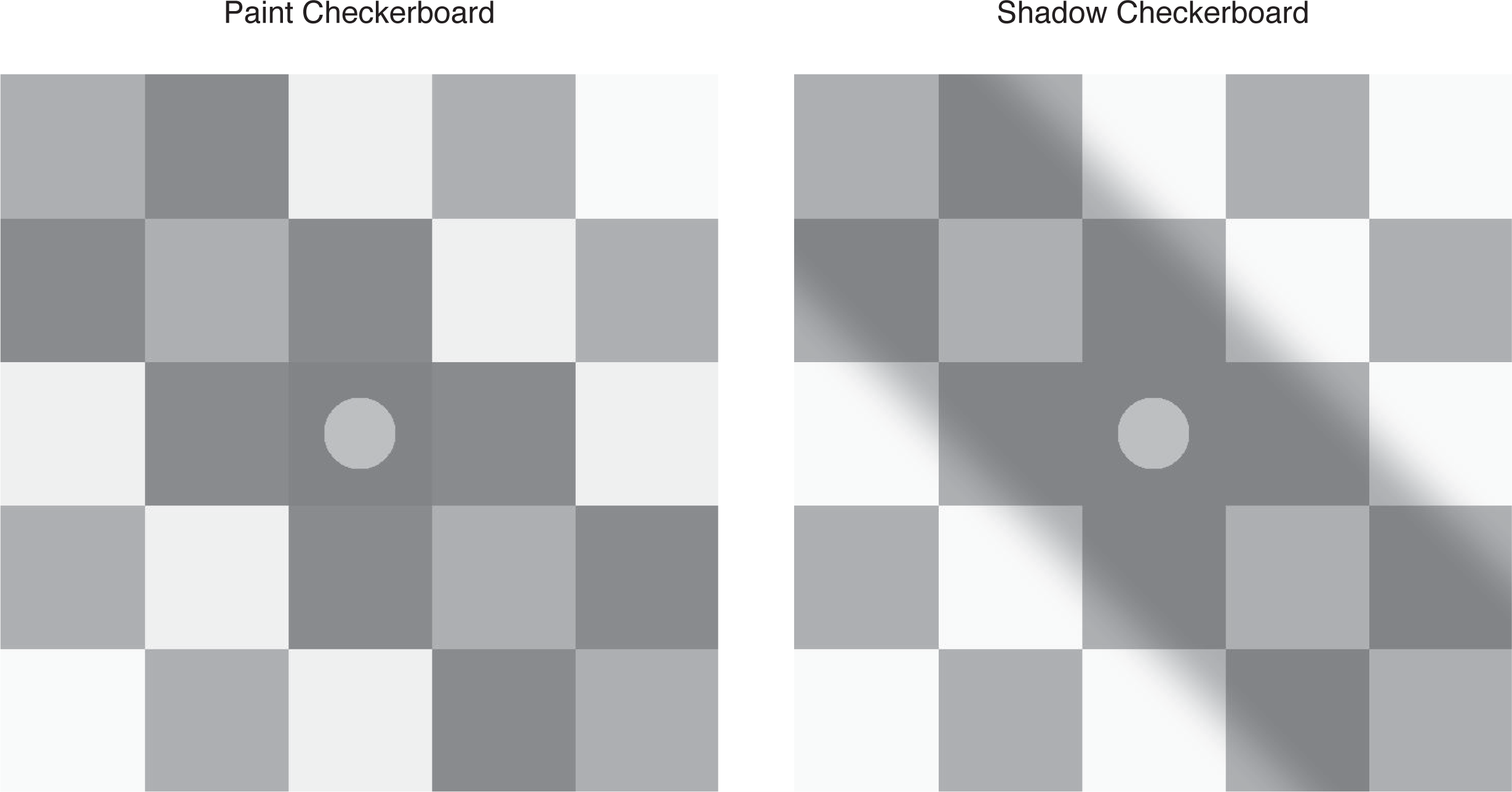
Paint and shadow checkerboard stimuli. The paint checkerboard is shown on the left and the shadow checkerboard on the right. The mean luminance of each corresponding square in the two checkerboards is identical, as is the luminance of the disks. Nevertheless, the disk in the shadow checkerboard appears lighter. This effect demonstrates the paint-shadow illusion for our stimuli. The disks illustrated have a luminance of 0.5.

The goal of this study was to understand the relationship between neuronal population responses and lightness perception. Previous studies in animal models have found that a small proportion of neurons in visual cortex respond in ways that are consistent with lightness perception in humans, but the responses of single neurons are heterogeneous (Rossi and Paradiso, 1996; Rossi et al., 1996; MacEvoy et al., 1998; Rossi and Paradiso, 1999; Kinoshita and Komatsu, 2001; MacEvoy and Paradiso, 2001; Roe et al., 2005; Vladusich et al., 2006; Hung et al., 2007; Huang and Paradiso, 2008). However, perception of complex stimuli is presumably driven by the joint activity of neuronal populations, and it is not currently clear how population neuronal activity is integrated to give rise to lightness perception. Functional imaging studies (e.g. using fMRI) that measure the BOLD signal, which reflects the integrated activity of many neurons, have reported varied findings regarding neural correlates of lightness (Haynes et al., 2004; Perna et al., 2005; Cornelissen et al., 2006; Boyaci et al., 2007; Pereverzeva and Murray, 2008; Corney et al., 2009; Boyaci et al., 2010).

Here we use a combination of human psychophysics, simultaneous recordings from dozens of neurons in areas V1 and V4 in monkeys, and neuronal population data analysis techniques to explore the way that luminance/contrast and context are encoded and might be read out. We found that neuronal populations in both V1 and V4 represent variation in both luminance and context. The relationship between the representations of luminance and context is complicated. We show that lightness information can be read out from the responses of neuronal populations in V1 and V4 in a manner that is consistent with the illusion illustrated in Figure 1, perhaps by neurons in premotor areas in parietal or frontal cortex that are thought to be involved in the formation of perceptual decisions (Gold and Shadlen, 2007; Heekeren et al., 2008). At the same time, we show that the neural representations in V1 and V4 do not obligatorily lead to lightness representations consistent with the illusion. Rather, the interpretation of the information in our recorded populations depends on how that information is read out. More generally, our work shows how analyzing the responses of neuronal populations as a whole can illuminate the neuronal mechanisms underlying perceptual and cognitive processes.

## Materials and Methods

### Visual stimuli

To better understand the neuronal and psychophysical underpinnings of lightness perception, we studied visual stimuli of the type illustrated in Figure 1. A central disk was embedded in a checkerboard image, either within a shadowed region (‘shadow’ checkerboard) or within a luminance-matched region without a shadow (‘paint’ checkerboard). We studied how the perceived lightness of the center disk depends on context, which in our study refers to the difference between the paint and shadow surrounding checkerboards. Our stimuli differ from the original checker-shadow illusion in that the lightness effect occurs across disks viewed in the center of two separate images, rather than within a single image.

Importantly, in our stimuli, the disks always had the same immediate surround (the center check is the same luminance in the paint and shadow versions) and the same average global surround. Indeed, the average luminance of each of the 25 corresponding checks in the paint and shadow checkerboards were the same. The only difference between shadow and paint checkerboards was the spatial distribution of light in the first and second off diagonals. In the paint version, each check was spatially uniform. In the shadow version, the luminance was governed by a cumulative normal computed as a function of the distance from the off-diagonals. This produced a penumbra-like gradient. For our stimuli, the paint-shadow illusion cannot be mediated by changes in local contrast nor by light adaptation to the overall luminance of the context images, because these two factors are matched. Indeed, disk luminance and disk contrast are perfectly correlated for our stimuli, so that our experiments do not distinguish between luminance and contrast representations. We reasoned that by silencing contrast and light adaptation, our stimuli would be more likely to reveal the action of cortical computations that support lightness perception (see Hillis and Brainard, 2007). For simplicity in the following, we will describe the disks in terms of their luminance; the reader should bear in mind that given our stimuli, we could have equally well have used a contrast representation with all else remaining unchanged.

The disk luminance values were expressed in normalized units that vary between 0 and 1. A luminance of 1 corresponded to about 260 or 300 cd/m^2^ for the psychophysical experiments (varying across the two monitors used, see below) and about 105 cd/m^2^ for the physiological measurements, with the exact value in each type of experiment varying as the monitors aged. In our normalized units, the mean luminance of each image was 0.485, and the check that immediately surrounded the disks had a luminance of 0.170.

Note that the paint and shadow stimuli themselves differed sufficiently, based on the spatial distribution of light in the first and second off diagonals alone, that both human and non-human primate subjects would almost certainly be able to reliably discriminate between the two.

### Psychophysical experiments

All human psychophysical procedures were approved by the Institutional Review Board of the University of Pennsylvania and the experiments were conducted in accord with the tenets of the Declaration of Helsinki.

Stimuli were displayed on one of two NEC Spectra View LCD monitors (Model PA241W, 24” display, maximum luminance 300 cd/m^2^, stimulus chromaticity [0.31 0.33]); PA271W, 27” display, maximum luminance 260 cd/m^2^, stimulus chromaticity [0.31 0.32]) from a distance of 57 cm using a pixel resolution of 1920 by 1200 pixels. The displays were controlled with 8-bit precision per channel using Matlab (The MathWorks, Inc.), with a combination of routines from mgl (http://gru.stanford.edu/doku.php/mgl/overview) and the Psychophysics Toolbox (psychtoolbox.org; Brainard, 1997; Pelli, 1997). The monitors were calibrated using standard methods (Brainard et al., 2002) and the non-linear input-output relation of each monitor channel was corrected using table lookup. Stimulus size in pixels was adjusted across the two monitors so that the visual angle of the stimuli was the same on each.

On each trial, human subjects viewed either two paint checkerboards side-by-side, or one paint and one shadow checkerboard side-by-side. In the latter case, the left-right position of the paint and shadow checkerboards was randomized across trials. Each checkerboard contained a disk in its central square (as in Figure 1). We refer to one disk as the reference disk and the other as the test disk. The left-right location of the reference disk was randomized across trials. There were three reference disk luminances, 0.25, 0.5, and 0.75 on the normalized [0-1] luminance scale, and across trials the reference disk appeared in each of the two types of checkerboards. The test disk’s luminance was adjusted over trials using staircase procedures. The subject’s task on each trial was to indicate which of the two disks appeared lighter. No complex elaboration about what was meant by the term “lighter” was provided to the subjects, and none reported to us that they found the term confusing. There is a large literature on the effect of instructions in judgments of lightness and color, see Radonjić and Brainard (2016) for a recent treatment.

The size of the checkerboards was 3.5° of visual angle, and they were presented centered vertically and with their centers located at +/− 3.5° horizontally. Each check in the checkerboards was a 0.7° square. The disks had a diameter of 0.35°, and were centered on the center square of the checkerboards.

Trials for each choice of checkerboard pairings (paint-shadow and paint-paint) were run in separate sessions. The paint-paint checkerboard pairing served primarily as a control. There were two separate staircases in each session for each choice of which checkerboard contained the reference disk and reference disk luminance, so 12 staircases per session in all (2 choices of which checkerboard contained the reference x 3 reference luminances x 2 staircases). For each combination of checkerboard containing the reference and reference disk luminance, one staircase was 2 up 1 down and the other was 1 up 2 down. At the start of each session there were 5 practice trials, chosen randomly from the set of possible trial types. The practice trials were followed by 20 blocks of 12 trials, where each block contained one trial from each of the staircases presented in random order.

Each individual staircase was thus 20 trials long – that is there were a fixed number of trials per staircase. Thus, there were 245 trials per session, including the 5 practice trials.

Four subjects (3 female, one male; ages 19-50) participated in the experiment. All were naïve as to the purpose of the study and had visual acuity of 20/40 or better as tested with a Snellen eye chart. The subjects ran two sessions of each condition (paint-shadow and paint-paint) to complete what we refer to as a single *determination* of the psychophysical paint-shadow effect. We made one such determination for subjects BAF and EJE, and two each for subjects AQR and CNJ. In cases where we aggregate data across subjects, we treat each determination separately, so that data from AQR and CNJ are weighted more heavily than those from BAF and EJE.

For the paint-shadow data, for each combination of reference disk luminance and location (reference in paint or shadow checkerboard), we combined the data from the two within-session staircases and fit a cumulative normal to them, using a maximum likelihood fitting method. The fit was implemented using the Palamedes toolbox (Kingdom and Prins, 2010; www.palamedestoolbox.org). The point of subjective equality (PSE; luminance corresponding to 50% lighter judgments) for each session was obtained from this fit. Thus, there were six paint-shadow PSEs obtained per session. These were aggregated across the two sessions for each subject/determination to find a paint-shadow effect, as explained in Results below. The same data analysis procedure was applied for the paint-paint data, although in this case the reference was always in a paint checkerboard. None-the-less, we gave each of the two paint checkerboards a nominal label to allow a parallel analysis, and obtained six paint-paint PSEs for each session as well.

### Electrophysiology experiments

#### Stimulus presentation, subjects, and electrophysiological recordings

Non-human primate subjects passively fixated while we presented single checkerboard stimuli and recorded neuronal responses. We presented visual stimuli on a calibrated CRT monitor (calibrated to linearize intensity, 1,024 × 768 pixels, 120-Hz refresh rate) placed 57 cm from the animal. We monitored eye position using an infrared eye tracker (Eyelink 1000, SR Research). We used custom software (written in Matlab using the Psychophysics Toolbox; Brainard, 1997; Pelli, 1997) to present stimuli and monitor behavior. We recorded eye position (1,000 samples per second), neuronal responses (30,000 samples per second) and the signal from a photodiode to align neuronal responses to stimulus presentation times (30,000 samples per second) using hardware from Ripple Microsystems.

The subjects in our physiological experiments were four adult male rhesus monkeys (BR, JD, ST, SY, *Macaca mulatta*, 8.8, 10.0, 9.0, and 9.3 kg, respectively). All animal procedures were approved by the Institutional Animal Care and Use Committees of the University of Pittsburgh and Carnegie Mellon University. Before training, we implanted each animal with a titanium head holder. Then, the animal was trained to passively fixate while we presented peripheral visual stimuli. Monkeys BR, JD, and ST were also trained to perform other visually-guided tasks that were not used in the current experiments. Once training was complete, we implanted a microelectrode array (Blackrock Microsystems). In monkeys BR and ST, we implanted a 10 x 10 microelectrode array in area V1. In monkeys SY and JD, we implanted a pair of 6 × 8 microelectrode arrays in V4. In monkey SY, both arrays were in V4 in the right hemisphere while monkey JD received bilateral V4 implants. We identified areas V1 and V4 using stereotactic coordinates and by visually inspecting the sulci. We placed the V1 arrays posterior to the border between V1 and V2 and placed the V4 arrays between the lunate and the superior temporal sulci. The two V4 arrays were connected to a single percutaneous connector.

The distance between adjacent electrodes was 400 μm, and each electrode was 1 mm long. At this depth, the electrodes were likely to be in the middle cortical layers, although the curvature of the brain relative to the array and other experimental factors make it difficult to be certain.

We recorded neuronal activity during daily experiments for several weeks in each animal. During each daily experiment, the monkeys were rewarded for passively fixating while we presented single checkerboard stimuli for 1000 ms. We varied the location, size and orientation of the stimuli across all of our experiments. The stimuli were generally positioned such that both the disk and at least some of the shadowed part of the shadow checkerboard (and corresponding region of the paint checkerboard) fell within the classical spatial receptive fields of the population of neurons we recorded, and typically spanned sizes between 4 and 15 degrees of visual angle. The stimuli were repositioned within and across days with the goal of changing the configuration of different parts of the image on the receptive field of different neurons. We define an experimental session as a set of all disk luminances in both paint and shadow contexts that were presented at one location, size, and orientation. Multiple sessions were often collected during a single daily experiment with different stimulus configurations randomly interleaved. Because V1 receptive fields are substantially smaller than those of V4 neurons, the stimuli typically covered a greater proportion of V1 surrounds but the sizes of stimuli were varied across experiments performed in both areas. We collected data that spanned luminance values between 0.05 and 1, in steps of 0.05, and analyzed data for stimuli where the disk was an increment relative to its immediate surround (disk luminance 0.20 or greater). Only trials where the monkey maintained good fixation were retained. Our data sets were limited by the availability of high quality neuronal recordings and the monkeys’ willingness to complete a sufficient number of behavioral trials. We included data from sessions where a) there were at least 5 trials for each intensity for paint and for shadow stimuli where the monkey maintained good fixation and b) where the best root-mean-squared decoding error of our linear population decoder variants (x-axis values on Figure 8B, C below) was less than or equal to 0.20. We adopted this latter criterion because it seemed most conservative to make statements about the relationship between the encoding of luminance and context in situations where luminance was encoded with reasonable fidelity. The RMSE criterion led to us exclude 4 of 6 sessions from monkey BR, 1 of 17 sessions from monkey ST, 17 of 28 sessions from monkey JD, and 2 of 128 sessions from monkey SY. Note that simply decoding each disk luminance with a null model that estimates all disk luminances as the mean disk luminance leads to a mean (across sessions) RMSE of 0.24.^1^ We verified that relaxing the RMSE exclusion criterion to this null model value of 0.24, which excluded only 5 of 179 sessions, did not affect our conclusions. The average number of total trials per session in the data set was 648. Our analyzed data set includes 18 recording sessions in V1 (2 from Monkey BR and 16 from Monkey ST) and 137 recording sessions in V4 (11 from Monkey JD and 126 from Monkey SY). Trial-by-trial data for each session, in the form of disk luminance, stimulus type (paint or shadow) and resulting spikes per electrode, are available in a public data repository at URL https://figshare.com/articles/Individual_Session_Data/5948077/1. The data can also be provided in a rawer form upon request.

**Figure 2.**
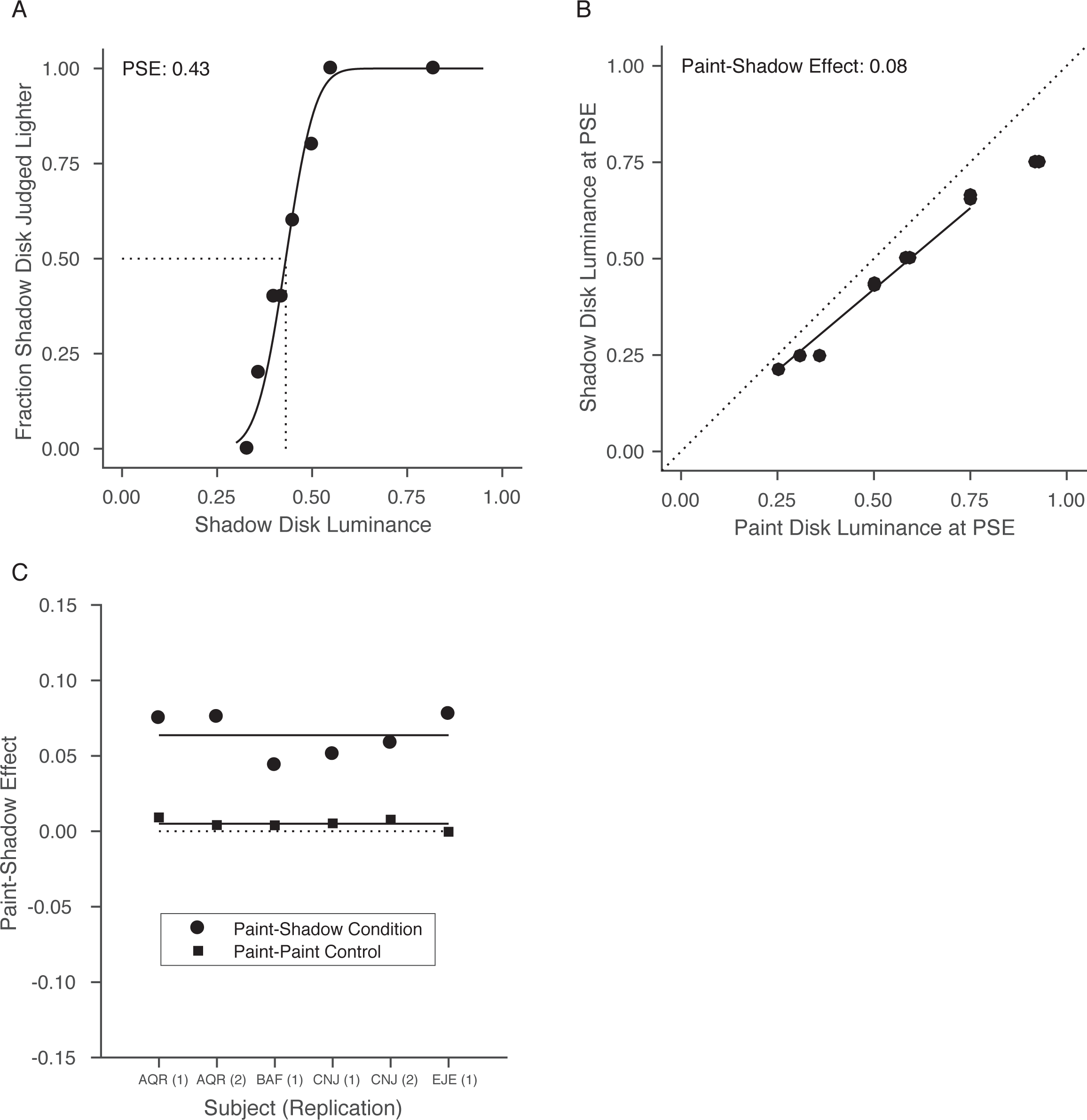
Human psychophysical experiments quantify the paint-shadow effect. **A**. Results from an example psychophysical session for one subject (AQR). The fraction of times that the test disk (in the shadow context) was judged lighter than the reference (in the paint context) is plotted as a function of test disk luminance. The reference disk luminance was 0.5. To produce the plotted points, test luminances generated by the staircase procedure were sorted and aggregated into groups of trials; the black line through the plotted points is the maximum likelihood cumulative normal fit to the individual-trial data. The PSE obtained from the fit is shown by the dashed line. The data quantify the observation that disks appear lighter in the shadow checkerboard than in the paint checkerboard (see Figure 1), as the luminance of the PSE is less than that of the reference. **B.** Paint-shadow effect for one subject (AQR) for data aggregated across two sessions. Each point represents a pair of disk luminances that match in appearance when one is presented in the paint context (x-axis) and the other in the shadow context (y-axis). For trials where the reference was in the paint context, the PSE is plotted against test luminance. For these trials, the PSE is generally less than that of the test, as in the case shown in A. For trials where the reference disk was in the shadow context, the test disk luminance is plotted against the PSE. This reversal keeps the sign of the effect shown in the figure consistent across the two types of trials. The paint-shadow effect is taken as the log_10_ of the slope of the best fit line to the data, with the line constrained to pass through the origin (best fit line shown). For these data, the slope is 0.84 corresponding to a paint shadow effect of 0.08. This indicates that the disks of the same luminance appear lighter in the shadow context than in the paint context. The line was fit using a least-squares criterion with the fit restricted to points on the x-axis in the range between 0.25 and 0.75. **C.** Summary of psychophysical paint-shadow effect. For each subject/determination, the paint-shadow effect, taken as the negative log_10_ of the slope, is shown as a solid circle. Across subjects/determinations, the mean paint-shadow effect is 0.06. For comparison, paint-shadow effects obtained from control conditions where both disks were presented in the paint context are shown as solid squares. As expected, these lie close to 0 (mean value of 0.005).

**Figure 3.**
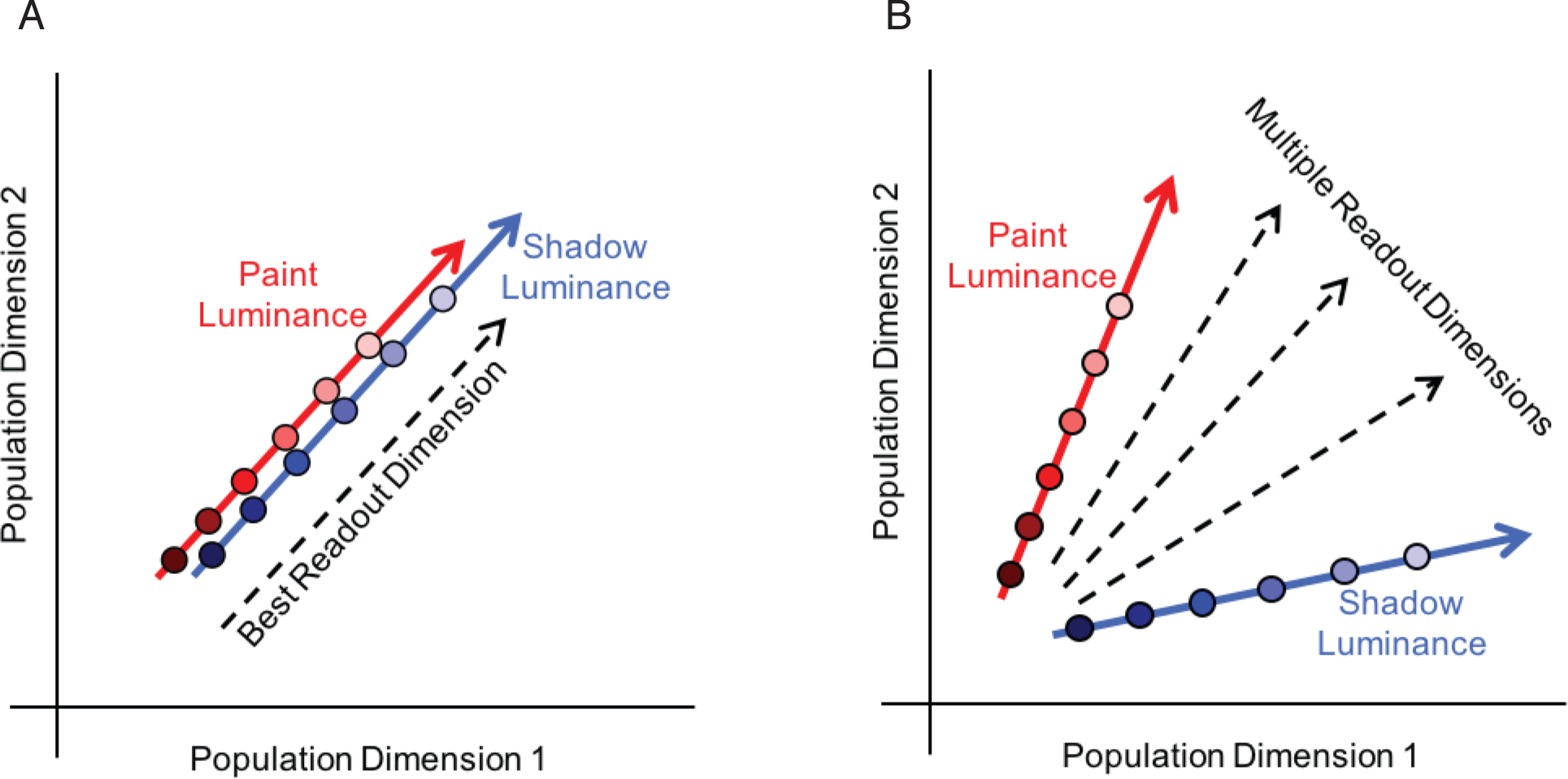
Schematic of candidate population neuronal mechanisms underlying the paint-shadow effect. These schematics depict two possible neuronal representations that could fulfill the criteria for an underpinning of the paint-shadow effect. Population activity in response to a single luminance/context condition is represented in each schematic as a single point in a neuronal population space. Each dimension in this space may be thought of as representing the firing rate of one of the simultaneously recorded neurons. In both schematics, the luminance of paint stimuli is encoded along the direction represented by the red arrow and the luminance of shadow stimuli is encoded along the direction represented by the blue arrow. **A.** Population activity for paint and shadow stimuli lies along a common locus in the same neuronal subspace, as disk luminance is varied. The best linear decoding dimension for stimulus luminance is illustrated by the dashed black line. In this case, luminance decoding for disks in shadow would be higher than that for disks in paint to the degree that the response for disks in shadow are shifted upwards along the common direction of variation in neuronal response space, and this neuronal paint-shadow effect would be observed for any reasonable luminance readout direction. **B.** Population activity for paint and shadow stimuli varies along separate directions. Decoding luminance by projecting the neuronal response onto any of the dimensions represented by the dashed black lines would give high fidelity luminance information that is modulated by paint-shadow context, with the degree and direction of the effect of the resulting paint-shadow effect determined by the choice of decoding dimension as well as exactly where responses for each disk luminance fell along the paint and shadow response directions (red and blue lines respectively).

**Figure 4.**
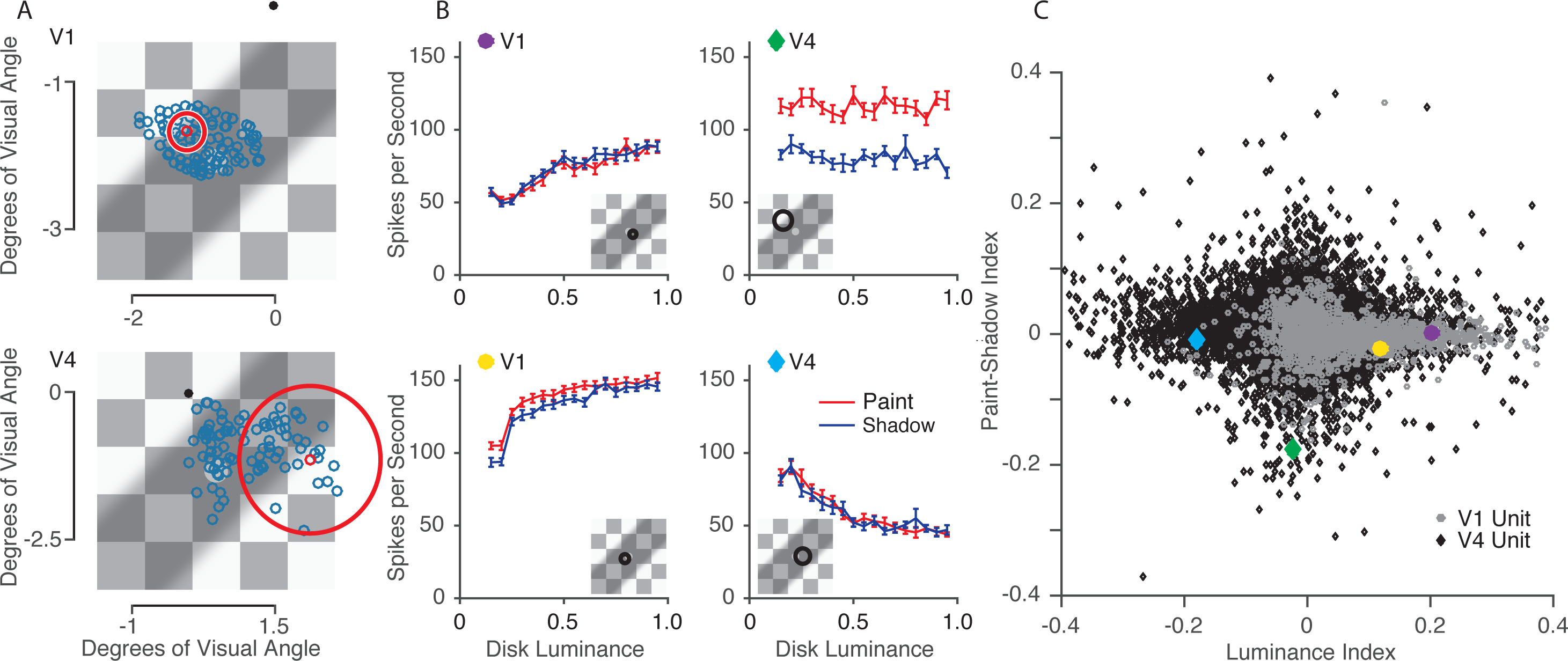
Responses to paint and shadow checkerboard stimuli are heterogeneous across the population. **A.** Example receptive field (RF) locations from a single session of V1 data (top) and V4 data (bottom) overlaid on an example stimulus. RF center locations were estimated by flashing small Gabor stimuli in a grid of positions while the monkey was rewarded for passively fixating. The black dot represents the fixation spot during the checkerboard stimulus presentation. The blue circles represent estimated RF center locations. The large red circle represents the estimated size of one example unit’s RF, with the RF center location drawn as the small red circle. Across sessions, the size, position and rotation of the checkerboard stimuli were varied. **B.** Luminance response plots for 4 example units. The brain area of these example units is denoted by the shape of the colored insets (circles: V1; diamonds: V4). An estimate of the size and location of each unit’s receptive field, relative to the checkerboard stimulus, is shown in the inset. **C.** Scatter plot of paint-shadow indices versus luminance indices for all V1 and V4 units (grey circles, 1,744 units; black diamonds, 11,063 units, respectively). The paint-shadow index was calculated using averaged responses to all disk luminances sorted by whether the stimulus was from the paint or shadow set, using the equation (shadow-paint) / (shadow+paint). Median population value for V1 = 0.0015, V4 = 0.0049. The luminance index was calculated using averaged responses from only the paint trials with disk luminances at 0.25 and 0.75 using the equation (resp75-resp25) / (resp75+resp25). Median population value for V1 = 0.037, V4 = −0.010.

**Figure 5.**
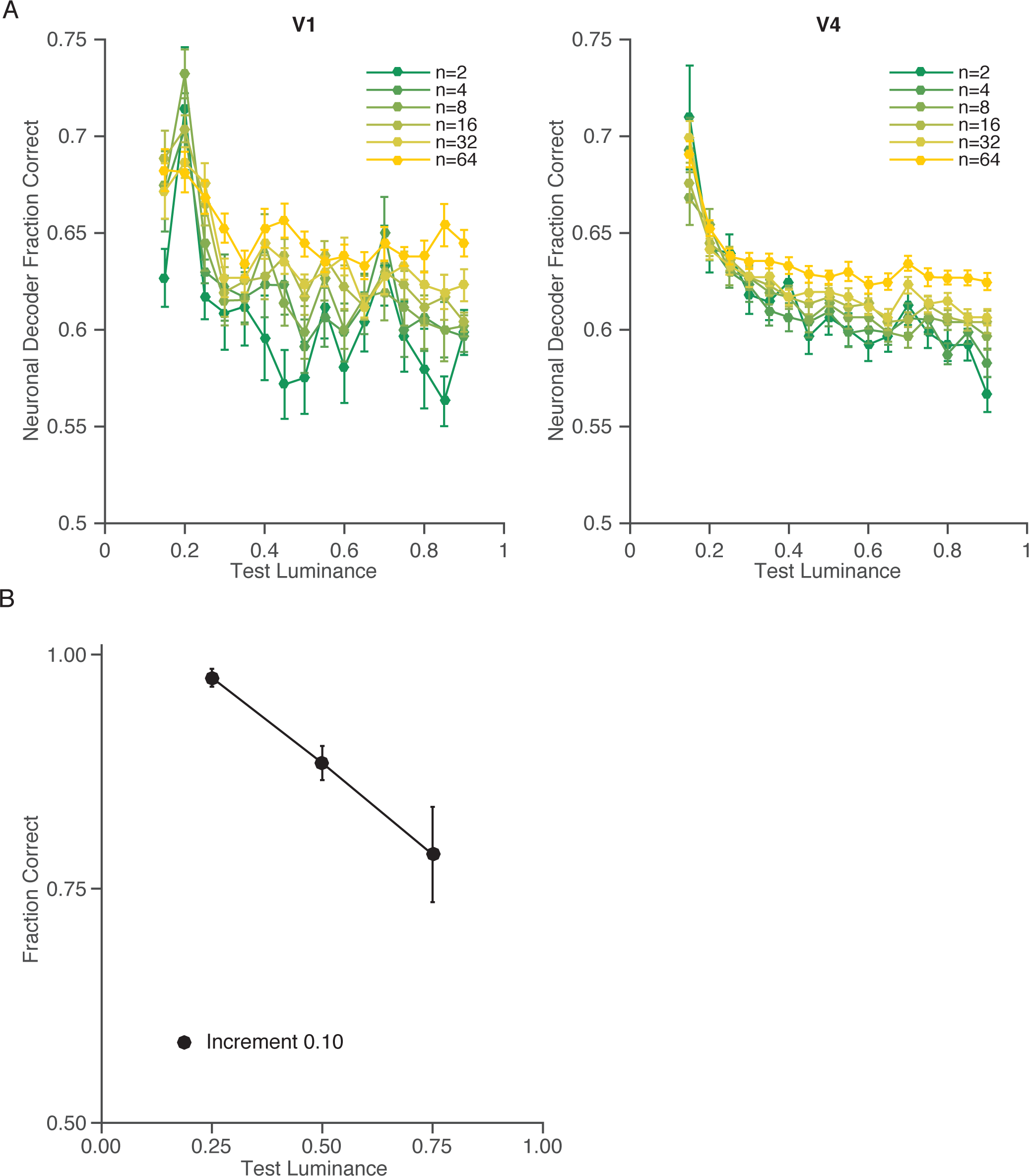
Neuronal and psychophysical sensitivity show similar dependence on reference luminance. **A.** Average precision of population encoding of luminance as a function of number of units, for each reference luminance for V1 (left, 18 sessions) and V4 data (right, 137 sessions). Performance of a neuronal decoder at discriminating 0.10 luminance increments as a function of the reference luminance and the number of units included. Data points represent the mean across all paint trials for data sets from V1 and V4 for 10,000 random draws per base luminance and population size, per data set. Error bars are the standard error of the mean across data sets. **B.** For the conditions where both disks were presented in the paint checkerboard, this plot shows the average (across subjects/determinations) probability of correctly judging a luminance increment of 0.10 as lighter, for the three reference luminances (0.25, 0.50, 0.75) used in the psychophysical studies. These data are taken from the paint-paint checkerboard pairings. Error bars show +/- 1 SEM taken across subjects/determinations (n=6, see Figure 2C).

**Figure 6.**
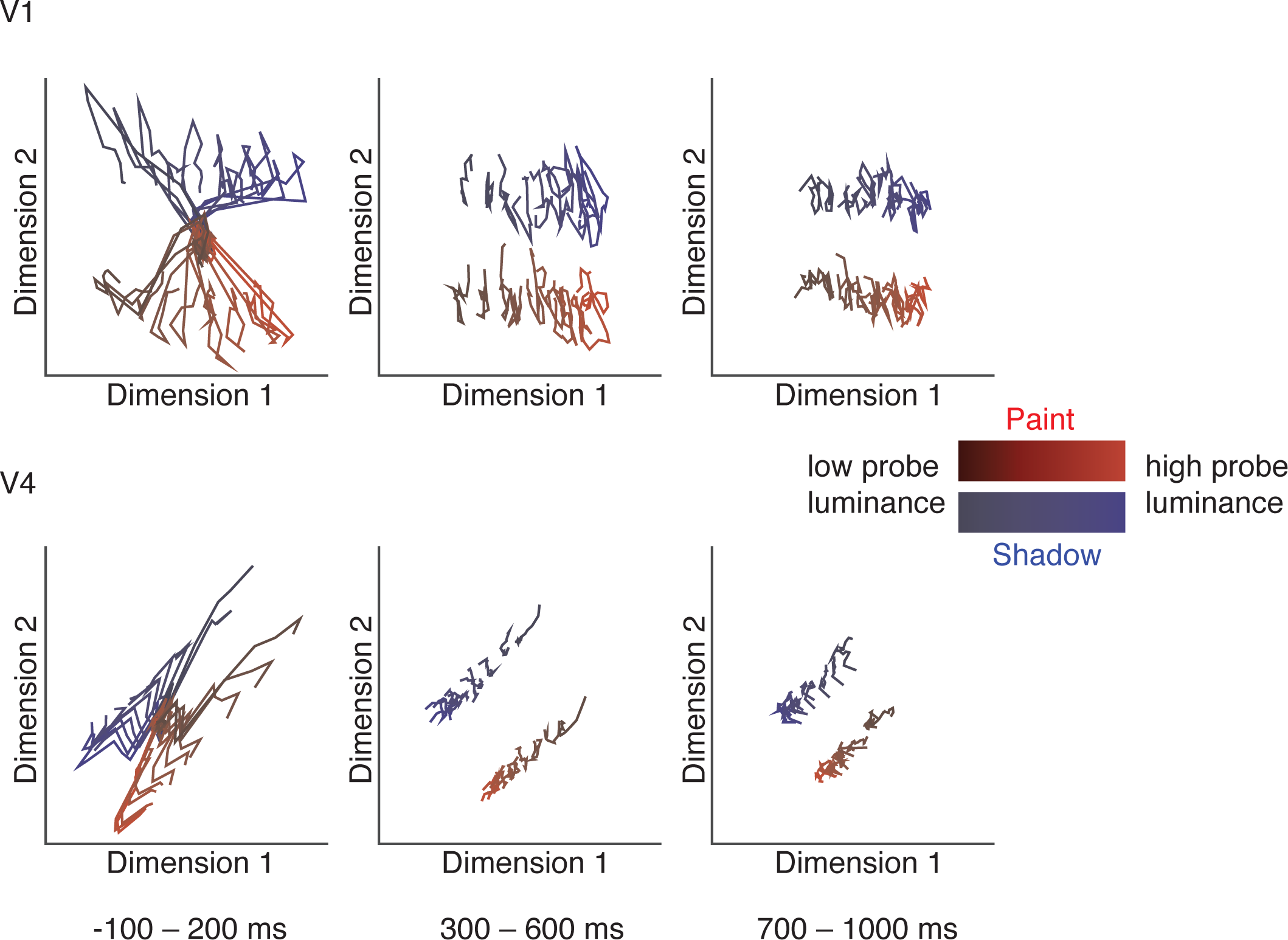
Population response trajectories for both V1 and V4 reveal consistent separation of responses for both context and disk luminance across the response period. Averaged responses from all conditions were combined across sessions for all recorded units. Gaussian Process Factor Analysis was used (20ms bin size) to identify the dimensions of population activity that explained the most variance in population responses. Linear discriminant analysis was used to select the single projection that best separates the all stimulus conditions for each area. These plots show the trajectories (averaged over all trials and all data sets, 18 sessions for V1, 137 sessions for V4) in each brain area for each of the luminances ranging between 0.25 and 0.75, and context during the beginning (left), middle, and end (right) of each stimulus presentation.

**Figure 7.**
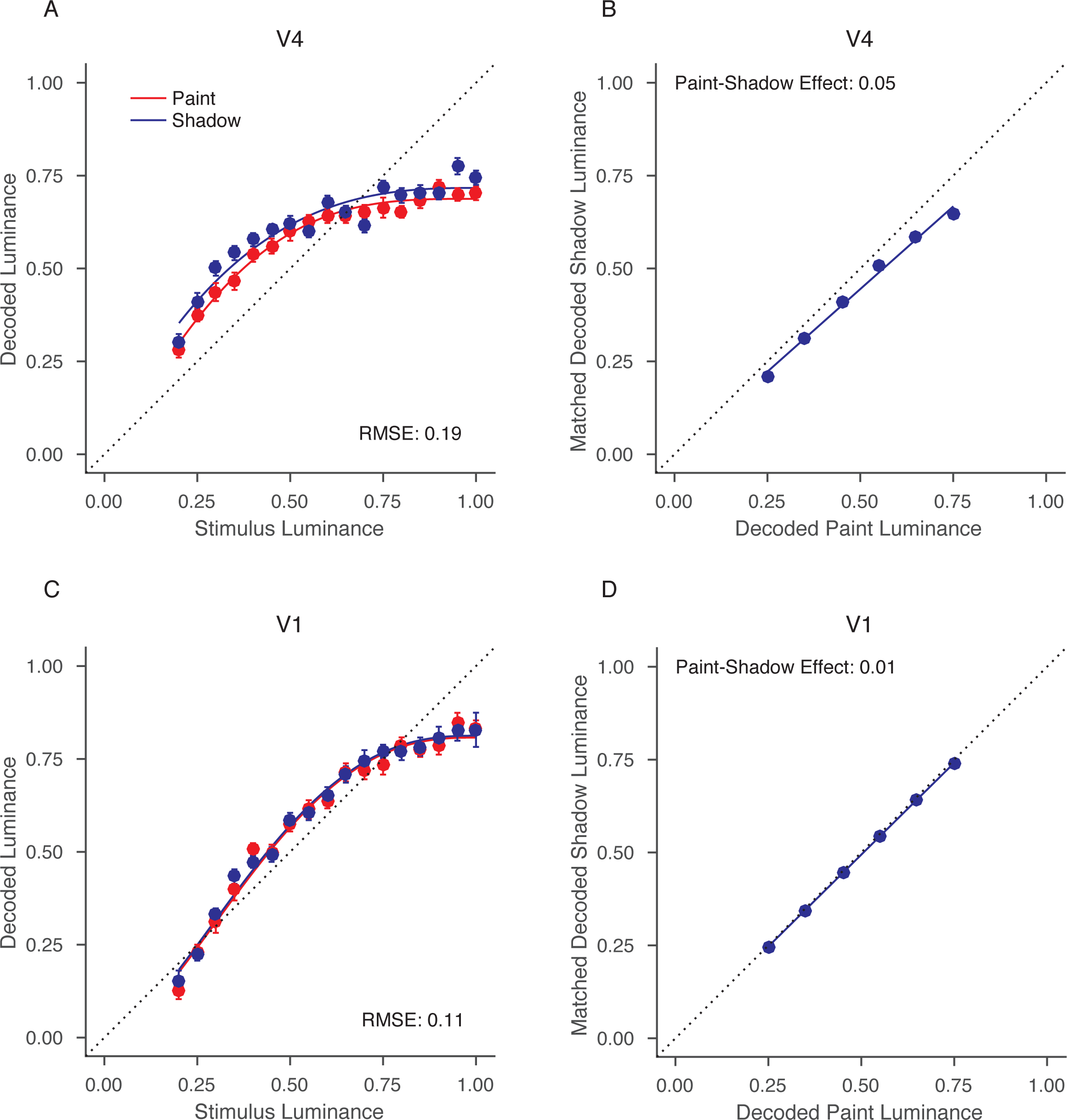
Computing a neuronal paint-shadow effect. **A.** Decoded luminance for paint and shadow stimuli, from a single session (JD, V4). The x-axis shows the stimulus luminance while the y-axis shows the decoded luminance. A single decoder was constructed to minimize RMSE for both stimulus types, with the results plotted separately for paint (red) and shadow (blue). Error bars show +/- 1 SEM and are often smaller than the plotted points. The smooth curves through the data are a fit affine scaling of the cumulative distribution of the beta probability density, a functional form chosen for convenience and not for theoretical significance. **B.** Paint-shadow effect derived from the decodings shown in A. Using the smooth fits to the decoded luminance, we found the disk luminances in shadow that were decoded to the same luminances as disk luminances in paint of 0.25 0.35, 0.45, 0.55, and 0.65, and 0.75. These disk luminances are plotted, with decoded paint luminance on the x-axis and matched decoded shadow luminance on the y-axis. A line through the origin was fit through these points and the negative log_10_ of the slope of this line taken as the paint-shadow effect, 0.05, for the decodings shown in A. **C.** Same as in A but for a session from a different monkey and visual area (ST, V1). **D.** Same as in B, for the decoding shown in C. Here the paint-shadow effect is −0.01.

**Figure 8.**
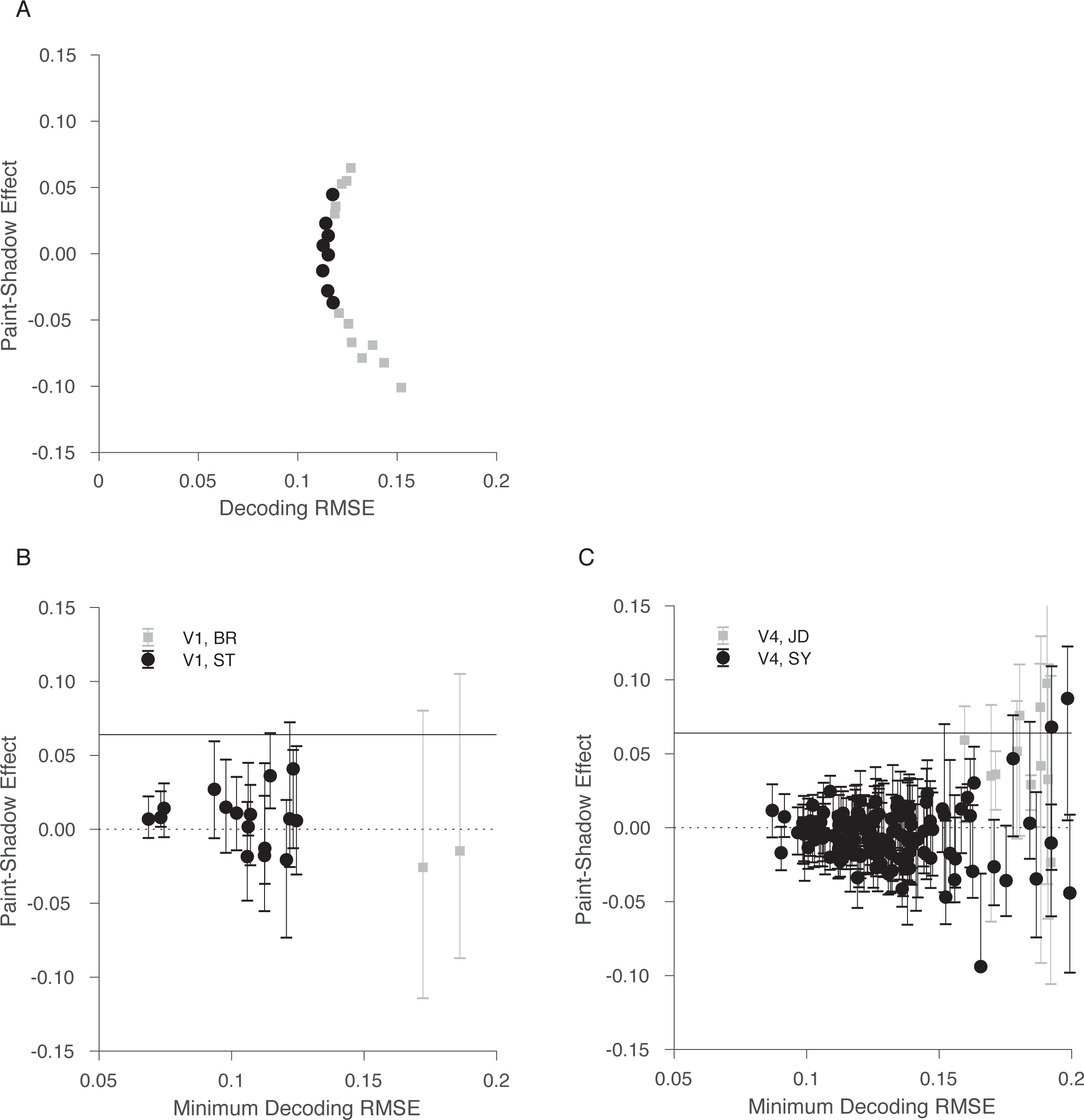
Decoding reveals large range of possible paint-shadow effects. **A.** Single V4 session example of paint-shadow effect as a function of decoding RMSE. The points shown as black circles represent cases where the decoding RMSE is less than 5% larger than its minimum value. We took as the plausible range of paint-shadow effects for this session the range corresponding to the black points. **B.** Summary across all included V1 sessions (n=18) of the range of obtainable neuronal paint-shadow effects as a function of decoding RMSE for each session. For each session, the ordinate of the plotted point is the paint-shadow effect from the decoding that had minimum decoding RMSE, while the abscissa is that minimum decoding RMSE (obtained for each session across the set of decoders examined for that session). The range bars extending from each point show the range of paint-shadow effects corresponding to the decoders where the RMSE was no more than 5% greater than the minimum. The psychophysically-determined paint-shadow effect is denoted by the solid black horizontal line. The dashed black horizontal line indicates a value of 0. Black circles: monkey ST; gray squares: monkey BR. **C.** Summary across all included V4 sessions (n=137), same format as B. Black circles: monkey SY; gray squares: monkey JD.

### Experimental Design and Statistical Analysis

All spike sorting was done manually following the experiment using Plexon’s Offline Sorter. We sorted single units as well as multi-unit clusters (multi-unit clusters, which comprise the majority of our data set, were sorted to remove noise). During recordings from the chronically implanted microelectrode arrays we used in V1 and V4, it was nearly impossible to tell whether we recorded from the same single- or multi-unit clusters on the array across subsequent days. Because of this, our primary analyses are based on neurons that were recorded simultaneously during a single recording session. For this study, we have combined data from single units and multi-units, and we use the term “unit” to refer to either. We included units for analysis if their response to the checkerboard stimuli was significantly different from the baseline response 100 ms prior to stimulus onset (t-test, p < 0.01, with Bonferroni correction for the number of units recorded during the session). We recorded from an average of 96 units per session from monkey BR, 97 from monkey ST, 55 from monkey JD and 83 from monkey SY. To allow for the latency of V1 and V4 responses, our analyses are based on spike counts calculated from 30-1030 ms after stimulus onset for V1 and 50–1050 ms after stimulus onset for V4. Figure 6 below analyzes response dynamics and indicates that our results are unlikely to be sensitive to the response interval chosen for analysis.

### Population decoding

To study how neuronal populations encode luminance and context, we used a linear decoding approach.

We used standard linear regression to predict disk luminance from the spike count responses of populations of simultaneously recorded units. We fit the luminance of paint and shadow trials (all together) as a linear combination of the responses of all simultaneously recorded units to stimuli with disk luminances greater than or equal to 0.2 (all stimuli where the disks were increments relative to their immediate surround). Specifically, we fit the model x = Yb + b_0_, where x is a #trials by 1 vector of disk luminances, Y is a #trials by #units matrix of spike count responses to the analyzed stimuli, b is a #units by 1 vector of weights, and b_0_ is a scalar affine term. We then assessed the decoded luminance of paint and shadow trials separately. To assess how well the luminance decoders performed, the root mean squared error (RMSE) of the luminance predictions was obtained by comparing the decoded luminances to the true disk luminances using 10-fold cross-validation.

We chose to study linear decoders because a) it is known that the computations required for linear decoding can be implemented by neurons and b) information that may be read out by a linear decoder is reasonably described as explicitly represented in the neuronal population (Pagan et al., 2013; Majaj et al., 2015). The latter point is important, as our primary interest is not in determining whether information about disk luminance and paint-shadow context is present in some form in V1 and V4, but rather in determining the degree to which this information explicitly supports a paint-shadow effect. In preliminary analyses, we also explored maximum likelihood decoders, which estimated disk luminance based on which luminance maximized the likelihood of the observed responses. These did not perform as well in a cross-validated RMSE sense as standard linear regressions, presumably because our data set does not contain enough trials to adequately estimate the unit response parameters (e.g., response mean, response variance) required to compute response likelihoods, and we do not report results based on maximum likelihood decoders. We also repeated our decoding analyses using trial-shuffled data to destroy noise correlations between simultaneously recorded units. Although correlated neurons contribute to the non-unique decoding weights we observed, destroying the correlation structure had a small effect on the decoding RMSE and did not change the key features of the paint-shadow effect shown in Figure 8B, C.

Note that our decoding methods allow any combination of weights on the units we recorded. If luminance information was best decoded using the responses of a small number of well-tuned units (perhaps those whose receptive fields best overlapped the stimuli we used), the weights of those units could be large while the weights on the rest of the population could be zero. In practice, however, the distributions of weights were broad, and the decodings were non-unique (see Results).

## Results

### Human subjects perceive higher lightness in the shadow than in the paint context

The main goal of the human psychophysical experiments was to quantify the paint-shadow effect for our stimuli.

An example psychometric function is shown in Figure 2A. This plots the fraction of trials on which test disks seen in the shadow checkerboard were judged to be lighter than the reference disk in the paint checkerboard, as a function of test disk luminance. The luminance of the reference disk was 0.5. Sensibly, as the test disk luminance increased it was judged lighter more of the time. The point of subjective equality (PSE) obtained from the maximum-likelihood cumulative normal fit to the data is indicated by the dashed line. This value was inferred from the fit and corresponds to the test disk luminance at which subjects would report that the test was lighter than the reference on 50% of trials. The difference between the PSE and the reference disk luminance shows the perceptual effect of context on lightness. Here, the luminance of the PSE is less than that of the reference disk, indicating in turn that disks of equal luminance appear lighter in the shadow checkerboard than in the paint checkerboard.

Figure 2B summarizes the psychophysical results for all of the paint-shadow measurements for the same subject/determination whose example psychometric function is shown in Figure 2A. Each point represents one PSE from a single session, with data from the two sessions of the single determination shown. For the cases where the reference disk was in the paint checkerboard, the reference disk luminance is on the x-axis and the PSE luminance is on the y-axis. For cases where the reference disk was in the shadow checkerboard, the PSE luminance is on the x-axis and the reference disk luminance is on the y-axis. In all cases, a lower luminance was required for a disk in shadow to appear the same as a disk in paint. We summarized this effect (which we refer to as the paint-shadow effect) as the negative log_10_ of the slope of the best-fit line through the PSE points (solid line in Figure 2B). The fit line was constrained to pass through the origin, and in fitting we only considered points where the x-axis value was in the range 0.25-0.75.^2^ This allows us to make a matched choice that avoids neuronal saturation when we perform a parallel analysis on the neuronal data below. For the data shown in Figure 2B, the slope was 0.84 and the paint-shadow effect was 0.08. The convention of choosing the negative, rather than positive, log_10_ slope makes a positive paint-shadow effect one that is consistent with the observation that disks of the same luminance appear lighter in shadow.

The results shown in Figure 2B were characteristic of the data we obtained from other subjects/determinations. Figure 2C plots the paint-shadow effect obtained for each subject/determination (solid circles). The mean effect was 0.064 (+/-0.006 SEM). We applied the same analysis procedures to control paint-paint checkerboard pairings (solid squares). Here, as expected, the mean paint-shadow effect was close to zero (0.005 +/− 0.001). The psychophysical data provide a quantitative measurement in humans of the magnitude of the paint-shadow illusion for our stimuli.

### Criteria for a neuronal explanation for the lightness illusion

The existence of lightness illusions and constancy tell us that the neuronal representation of lightness combines information about the luminance of light reflected from objects with information about the context in which they are viewed. The goal of the physiological part of this study is to understand how information from these two separate sources is represented in populations of cortical neurons, and in particular the degree and manner to which the neuronal representation of disk luminance is affected by variation between the paint and shadow contexts. Our guiding hypothesis is that luminance and context are represented jointly in neuronal populations in visual cortex, and that this ultimately leads to a context-dependent transformation of luminance to lightness as the information is read out by subsequent processing stages. More specifically, we test the idea that parsimonious accounts of decoding surface lightness from the measured population can provide a higher readout for disks in the shadow checkerboard than for disks in the paint checkerboard. If this is the case, the nature of the population code reveals a mechanism that can contribute to the visual system’s ultimate representation of surface lightness. Thus, we suggest that the way that disk luminance and context are encoded in a candidate neuronal population should satisfy the following criteria:

*Criterion 1*: Disk luminance should be encoded with good fidelity in ways that are broadly consistent with human discrimination psychophysics. For example, human subjects are more sensitive to subtle luminance changes of a disk when the luminance of the disk is low than when it is high.

*Criterion 2*: Context should also affect the population responses when disk luminance is held fixed, so that the readout of disk luminance could be affected by context.

*Criterion 3:* Plausible methods of reading out lightness from the population responses, regardless of whether the responses of individual neurons themselves vary monotonically with luminance, should accommodate a context effect in the same direction as the illusion, so that the read-out lightness of a disk in shadow is higher than that of a corresponding-luminance disk in paint. In addition, the same readout should account for the fact that lightness increases with luminance when context is held fixed.

To provide intuition for how neuronal representations could fulfill the above criteria, Figures 3A and 3B illustrate two scenarios. The schematics in each panel show a neuronal population space. Each dimension in this space could be taken to represent the firing rate of one of the simultaneously recorded neurons (so a population of 100 neurons would be represented in a 100-dimensional space). The response to a visual stimulus would then be represented by a point that indicated the number of spikes each unit fired during the stimulus presentation. More generally, a neuronal population space could be a lower dimensional projection of the individual neuron firing rate space. In both schematics shown, the variation in neuronal response to changes in the luminance of disks in the paint checkerboard is encoded along the direction represented by the red arrow, while the luminance of disks in shadow checkerboards is encoded along the direction represented by the blue arrow.

In the scenario represented by Figure 3A, varying the luminance of disks in both paint and shadow causes the neuronal representation to vary along a single direction, with the effect of context being to shift the representation of disks in shadow along this direction relative to the representation of disks in paint. Here it would be natural to read out the lightness of the disk by projecting the responses onto a readout dimension that was aligned with the common direction of stimulus variation. This dimension is shown in Figure 3A by the dashed black arrow, with the position of the arrow shifted laterally from the red and blue arrows to avoid excessive clutter in the depiction. Because the effect of paint versus shadow context shown in 3A is to shift the representation of disk luminance along the single direction of variation, the lightness decoder illustrated will produce an obligate paint-shadow effect. This coding idea underlies, at least implicitly, a number of single-unit and fMRI studies of the neuronal representation of lightness, in which the question posed is whether the response magnitude of individual units or voxels to luminance is shifted by context in a direction consistent with perceptual effects evoked by the stimuli under study (Rossi and Paradiso, 1996; Rossi et al., 1996; Rossi and Paradiso, 1999; Kinoshita and Komatsu, 2001; MacEvoy and Paradiso, 2001; Haynes et al., 2004; Perna et al., 2005; Roe et al., 2005; Cornelissen et al., 2006; Boyaci et al., 2007; Pereverzeva and Murray, 2008; Boyaci et al., 2010).

In the scenario represented by Figure 3B, the direction of response variation corresponding to varying disk luminance is different for the paint and shadow contexts. As in 3A, the neuronal representation of luminance is affected by context, but in a qualitatively different manner. Here multiple possible readout directions for decoding lightness are shown (black dashed arrows), and across these there are potential tradeoffs between the precision with which the read-out lightness encodes within-context luminance variation and the degree to which the readout will reveal a paint-shadow effect. The goal of our study is to determine whether the representation of lightness in early visual cortex caries with it a requisite paint-shadow effect (as in Figure 3A) or whether, as in Figure 3B, there are many equivalent readout dimensions, which carry with them a range of paint-shadow effects.

Note that these schematics are simplified for illustrative purposes. They show the effect of luminance variation as lines in a one-dimensional subspace of a two-dimensional space, but the actual dimensionalities are higher. In addition, in higher dimensions, the variation may trace out non-linear paths within the subspaces they occupy, and the subspaces occupied by the paint and shadow could share some dimensions but diverge in others.

These considerations add richness beyond what is shown in the schematics. Below, we analyze our data to understand the relationship between lightness perception and the neuronal population representations of luminance and context.

### Neuronal populations in V1 and V4 encode luminance and context

Based on previous studies (Leopold and Logothetis, 1996; Sheinberg and Logothetis, 1997; Rossi and Paradiso, 1999), we chose primary visual cortex (V1) and V4 as areas in which to examine the neuronal representation of disk luminance and the effect of context. We recorded simultaneously from several dozen units in each area and positioned the stimuli so that they overlapped the receptive fields of the recorded units (Figure 4A). The exact positioning, size, and orientation of the checkerboard was varied across sessions (see Methods).

In both areas V1 and V4, we found individual units that were selective for luminance and/or context. Figure 4B shows the mean firing rates of four example units as a function of luminance in the paint (red) or shadow conditions (blue). The responses of these four example units are quite heterogeneous. For example, the unit shown in the upper left panel is modulated by disk luminance but there is little if any effect of paint versus shadow context. In contrast, the unit shown on the upper right is modulated by context but shows little modulation by disk luminance.

This heterogeneity was typical of our data set (Figure 4C, example units are specified by the colored inset in each plot in Figure 4B), and is expected given that we recorded from neurons with a range of receptive field locations and tuning properties. We characterized each unit by a luminance index and a paint-shadow index (see definitions in caption to Figure 4). A positive luminance index indicates that a unit responds to paint stimuli with a higher firing rate as disk luminance increases. A positive paint-shadow index indicates that a unit responds more to shadow stimuli than paint stimuli when disk luminance is equated, irrespective of whether this is a response to the central test disc itself or to any other component of the display. Thus luminance and paint-shadow indices of the same sign (first and third quadrants in 4C) indicate neurons whose individual response properties are consistent with the psychophysics, while indices of opposite sign (second and fourth quadrants) indicate neurons whose response properties go in the opposite direction from the psychophysics.

Consistent with reports in the literature of the existence of atypical luminance and contrast response tuning curves (Bushnell et al., 2011; Sani et al., 2013), our data set contained many units that had either positive or negative luminance or paint-shadow indices. In both areas and for both indices, the population means and medians were close to zero. Because it is difficult to determine the extent to which we recorded from the same units on subsequent days and thus the extent to which the data across days are independent, it is difficult to determine whether deviations from zero mean are statistically significant. In V1, the average luminance index was 0.072 (standard deviation = 0.091) and the average paint-shadow index was 0.0087 (standard deviation = 0.026). In V4, the average luminance index was −0.0093 (standard deviation = 0.082) and the average paint-shadow index was 0.0048 (standard deviation = 0.042). However, a central tendency single-number summary of the population response (e.g. mean or median) is not the most useful measure for connecting the neural measurements to perception. Rather, understanding the neural basis for (e.g.) the checker-shadow illusion requires understanding how the responses of many neurons (or a subset of neurons) may be read out to guide lightness perception.

The luminance and paint-shadow indices did not strongly depend on the relationship between their receptive field locations and the visual stimuli, and this relation did not explain a large proportion of the variance in the population data. The slopes of the best fit lines relating the absolute value of the luminance index and the distance between the center of the unit’s receptive field and the center of the disk were very close to 0 (--0.045 for V1 and 0.0015 for V4). Similarly, the slopes of the best fit lines relating the absolute value of the paint-shadow index and the distance between the receptive field center and the nearest diagonal line across the checkerboard that differentiates the paint from shadow stimuli were also very close to 0 (−0.0039 for V1 and 0.0017 for V4).

Despite these weak relationships, it remains likely that the way V1 and V4 neurons respond to visual stimuli depend on the way those stimuli fall on their receptive fields. Indeed, earlier single-unit studies, where stimuli were chosen with respect to the properties of individual neurons, none-the-less found modest contextual modulation of the activity of single cortical neurons in situations where context affects lightness, and observed considerable neuron-to-neuron heterogeneity (for review, see Paradiso et al., 2006). Our data set is more diverse, and we think more closely approximates the range of neuronal responses involved in lightness perception in natural vision, in which the heterogeneity of single neuron responses is presumably overcome by basing percepts and behaviors on the activity of large neuronal populations.

The heterogeneity of selectivity to luminance and context is reminiscent of the mixed and uncorrelated selectivity of neurons in many sensory areas to different stimulus features (for example, in primate area MT; DeAngelis and Uka, 2003; Smolyanskaya et al., 2013). Such mixed selectivity may be a natural consequence of a common underlying neuronal architecture as well as the fact that the neurons we recorded are tuned for many other stimulus features (e.g. orientation, spatial frequency, temporal frequency, color, texture, etc.). When selectivity is mixed, using the responses of many neurons, regardless of their specific tuning properties, may be advantageous for coding (Rigotti et al., 2013; Fusi et al., 2016).

We hypothesize that despite the heterogeneity we observe across units, the population response structure might support a parsimonious readout consistent with lightness perception. We next evaluate this hypothesis using the three criteria described above.

### Criterion 1: Sensitivity of the neuronal population to small changes in disk luminance

A basic prerequisite for a neuronal population to be part of the neuronal substrate for lightness perception is that it encodes luminance with reasonable precision. We asked how the neuronal population responses could be used to discriminate between a disk of particular luminance and one of increased luminance. Figure 5A shows the average ability of a cross-validated linear classifier to detect luminance increments of 0.1 as a function of the number of units (color) and the base disk luminance (x-axis) to which the increment was added for both V1 (left) and V4 data.

The classifier performs above chance for all choices of number of units, and performance worsens as base disk luminance rises. This is also true for the data from each individual animal (data not shown). The exact quantitative performance of the decoder depends on experimental factors such as the number of units we recorded from, the amount of noise in the recordings, the location of and size of the stimulus on the retina, and the extent to which the central disk overlapped with the receptive fields of the units. It is notable that, in addition to being above chance, the performance of the decoder shares a key feature with the psychophysical data. This feature is shown in Figure 5B, where we plot the average fraction of trials on which psychophysical subjects correctly judged that a disk with given base luminance plus an increment of 0.1 was lighter than a disk of the base luminance alone, when both disks were presented in the paint checkerboard. Here too we see a decrease in fraction correct with base luminance, a phenomenon that is generally referred to as Weber’s Law. The fact that neuronal population sensitivity matches, qualitatively, this feature of the psychophysics supports the idea that the neurons we recorded are involved in the processing that transforms luminance to lightness -- these neuronal populations satisfy the first criterion we proposed for a candidate neuronal explanation for the lightness illusion.

### Criterion 2: Context affects neuronal population responses

The schematics in Figure 3 depict two neuronal population representations of luminance that would lead to a paint-shadow effect: one in which the neuronal representations of paint and shadow overlap in population space and one in which the paint and shadow stimuli vary along different directions. To begin to differentiate between these possibilities, we assessed the similarity of the representations of paint and shadow stimuli by visualizing the mean responses in each context and luminance conditions as a function of time throughout the stimulus viewing period. To do so, we binned the responses in 20 ms bins, performed dimensionality reduction (using Gaussian Process Factor Analysis, (Yu et al., 2009; Cowley et al., 2013) https://users.ece.cmu.edu/~byronyu/software/DataHigh/datahigh.html), and plotted the trajectories for each luminance and context (snapshots of projections from the GPFA representation onto the best two dimensions, as determined by linear discriminant analysis, for each cortical area and several time bins are shown in Figure 6). This analysis was done using the entire data set, combining across sessions. It suggests that although there are interesting dynamics to the raw population responses, information about both luminance and context is represented in both V1 and V4 throughout the response period, and that there is no large qualitative change in the representation of either luminance or context across the response interval. For this latter reason, all of the other analyses in this paper are performed using data aggregated across the entire response interval (see Methods).

### Quantifying the neuronal paint-shadow effect

To relate the neuronal representations of disk luminance to the perceptual paint-shadow effect, we need a method to quantify the neuronal paint-shadow effect in a manner that allows comparison to the psychophysically-measured paint-shadow effect. Our approach was to use a linear regression technique to decode a neural correlate of disk lightness from the population responses, and then determine whether the result, which we refer to as the *decoded lightness,* differs for paint and shadow disks in a way that is consistent with the psychophysics.

To illustrate the idea, we begin by considering decoders that recover disk luminance from neuronal population responses. We used linear regression to predict disk luminance. We fit the luminance of paint and shadow trials (all together) as a linear combination of the responses of all simultaneously recorded units to stimuli with disk luminances greater than or equal to 0.2 (all stimuli where the disks are increments relative to their immediate surround). We then assessed the decoded luminance of paint and shadow trials separately. To assess how well the luminance decoders performed, the root mean squared error (RMSE) of the luminance predictions was obtained by comparing the decoded luminances to the true disk luminances using 10-fold cross-validation. Figure 7A shows the mean decoded luminance obtained in this manner for an example V4 session for paint disks (red) and shadow disks (blue), as a function of stimulus luminance. The RMSE for this example decoding was 0.19, which may be compared to a null model value of 0.24 that would be obtained from simply assigning to every disk the mean luminance of all of the presented disks. The improvement in decoding RMSE relative to the null value indicates that the units recorded in this session carried information about disk luminance. In this session, disks in shadow were decoded to higher luminances than disks in paint.

This suggests the possibility of using the luminance decoder as a way to generate decoded lightness. Indeed, for this example session, taking the decoded luminance as decoded lightness, the result is qualitatively consistent with the perceptual paint-shadow effect, where disks in shadow are perceived as lighter.

We examined whether the decoders tended to draw on the responses of just a few units, or more broadly on the responses of many units. For each session, we ordered the absolute value of the decoding weights, from largest to smallest. We then computed the sum of these weights, and asked how many units accounted for 25%, 50%, and 75% of that sum. We expressed these numbers as a percent of the total number of units for that session. For a decoder that draws primarily on just a few units, a small number of units would account for most of the absolute weight. We found that, on average, it required 5% of units to account for 25% (+/- 2% standard deviation) of the total absolute weight, 16% (+/- 5%) to account for 50% of the total, and 36% (+/- 6%) to account for 75%. Thus, on average, ~30% of the units contributed to the central 50% (25% to 75%) of the total absolute weight. We interpret this as indicating that the optimal decoder draws broadly on the responses of the neural population. As a check on this conclusion, we repeated the analysis using cross-validated lasso regression. Lasso regression uses an L1-norm regularization term to minimize the number of non-zero weights obtained in the regression solution, and thus provides a more conservative approach. We choose the regularization hyperparameter based on a five-fold cross-validation procedure, in which we evaluated the cross-validated RMSE error as a function of the regularization hyperparameter (25 values logarithmically spaced between 10^-5^ and 10) and chose the value that gave the lowest cross-validated RMSE. Using this value, we reanalyzed our dataset with lasso regression. We found weight distribution values very similar to those obtained with standard linear regression. (With lasso regression, it required 5% of units to account for 25% (+/- 2% standard deviation) of the total absolute weight, 15% (+/- 4%) to account for 50% of the total, and 34% (+/- 5%) to account for 75%. Thus, even with the more conservative lasso regression method ~29% of the units (on average) contributed to the central 50% (25% to 75%) of the total absolute weight.)

To quantify the neuronal paint-shadow effect for the illustrative decoding shown in Figure 7A, we fit the relationship between stimulus luminance and decoded luminance/lightness with a smooth function (fits shown as solid lines in 7A), and then used the fits to identify luminances for disks in paint and disks in shadow that decoded to the same value. Figure 7B plots pairs of *neuronal matches* (solid blue circles) obtained in this manner. We quantified the neuronal paint-shadow effect as the negative log_10_ of the slope of the best fitting line through the plotted points, with the line constrained to pass through the origin.^3^ For the data shown in 7B, the neuronal paint-shadow effect was 0.05. Figure 7C and D show the same analysis for an example V1 session, where the neuronal paint-shadow effect was found to be small (0.01).

A feature of the illustrative analysis shown in Figure 7 is that it reflects the combined effect of the way that paint and shadow stimuli are represented in the neuronal population and the action of the particular way we chose to build the decoder. That is, for illustrative purposes we equated decoded lightness with the output of a decoder built to estimate stimulus luminance. As depicted in the schematic shown in Figure 3B, when the population response to paint and shadow stimuli is multi-dimensional, there may be multiple decoders that can read out a neural correlate of lightness that preserves information about variation in disk luminance with high-fidelity. Indeed, the decoding approach we illustrated in Figure 7 was one that sought to find the same decoded luminance for both paint and shadow disks; that is, it was a decoder that sought to minimize the inferred neuronal paint-shadow effect. Because our interest is in whether the neuronal population codes can support the observed lightness effects, simply restricting the analysis to a decoder built to estimate luminance, as we did for illustrative purposes, is not appropriate. Rather, we want to characterize the range of paint-shadow effects that emerge when we explore a set of decoders, while at the same time requiring that the decoders preserve information about luminance variation.

### Criterion 3: The neuronal representations in V1 and V4 may be read out with high precision in a manner that produces a paint-shadow effect

To explore the range of neuronal paint-shadow effect obtainable with high-fidelity linear decoders, we determined the cost in decoding quality (quantified as root-mean-square error, RMSE) when we introduced a paint-shadow gain into the regression. That is, rather than constructing a single lightness decoder that attempts to estimate veridical disk luminance, we constructed a set of lightness decoders by introducing different estimation targets for paint disks and for shadow disks. We did this by defining a gain factor *g*, and dividing the target decoded paint luminance by *g* while multiplying the target decoded shadow luminance by *g*. The gain factor *g* implicitly sets a target paint-shadow effect for the decoder, and in this sense the decoders illustrated in Figure 7 had a target paint-shadow effect of zero. We varied g at 20 levels between 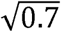 and 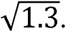. These correspond to paint-shadow effects between −0.15 and 0.11, a range that encompasses the human paint-shadow effect as well as an equal-sized effect in the opposite direction. For each resulting decoder we obtained the paint-shadow effect using the procedure illustrated by Figure 7. Figure 8A shows the obtained paint-shadow effects as a function of decoding RMSE for an example V4 session. Here the decoding RMSE was computed with respect to the target values, that is with respect to paint disk luminance divided by *g* and shadow disk luminance multiplied by *g*. We found that we obtain a wide range of paint-shadow effects (for this session between approximately −0.11 to 0.07) without a large effect on the decoding RMSE. If we restrict attention to RMSE values within 5% of the minimum value we found across all regressions, the range is still substantial (black filled circles in 8A), approximately −0.04 to 0.04.

Figure 8B, C shows for each experimental session the range of paint-shadow effects we obtained from lightness decoders whose RMSE was within 5% of the best RMSE obtained across all the examined decoders. The ranges were typically large (mean 0.077 in V1 and 0.054 in V4). The range straddles 0 for 81% of 155 recording sessions (89% of 18 V1 sessions, including both sessions from monkey BR and 14/16 sessions from monkey ST, and 80% of 137 V4 sessions, including 7/11 sessions from Monkey JD and 103/126 sessions from Monkey SY) and is strictly less than 0 for about the same number of sessions as it is strictly greater than 0 (8% of 155 sessions strictly positive compared with 11% strictly negative).

Our results show that many (although not most) of the high-precision lightness decodings lead to a paint-shadow effect that is in the direction of the psychophysics. These data are therefore consistent with the idea that the populations of V1 and V4 units we recorded satisfy the final criterion for a candidate neuronal explanation for the lightness illusion.

That said, the neuronal paint-shadow effect we observe is not obligate; one can construct high-precision decoders that do not show the paint-shadow effect as well as ones that do. In addition, the upper-edge of the decoded paint-shadow effect range is frequently smaller in magnitude than the mean psychophysical paint-shadow effect (see horizontal solid black line in Figure 8B, C). These observations suggest a population representation more like that depicted in Figure 3B than in 3A (see also Figure 6). Thus, the integration of context and luminance we observe in V1 and V4 neuronal populations provides a mechanism for computations that begin to extract perceptual lightness from stimulus luminance, but these populations do not reveal directly the full computations nor the specific readout mechanisms that would link the population code directly to the perceptual effects.

## Discussion

### A candidate neuronal explanation for perceived lightness

We combined human psychophysics, simultaneous recordings from dozens of neurons in V1 and V4, and neuronal population analyses to investigate the neuronal population mechanisms underlying lightness perception. With the psychophysics, we quantified a lightness illusion in the form of a measured paint-shadow effect. With neuronal recordings for the same stimuli, we found that the population representation of disk luminance is affected by the context in which the disk is presented. We found that the nature of the population representation allowed a range of high-precision decoders.

Although it was generally the case that this range included ones that produced neuronal paint-shadow effects in the direction consistent with the psychophysics (for which the neuronal lightness of disk of a given luminance was higher in the shadow than in the paint context), it was also the case that the range included ones that produced the opposite effect.

At first it might seem maladaptive to have context alter the lightness of disks that share the same luminance. Such contextual interactions, however, can be useful if we regard the function of lightness perception as providing a stable representation of object surface reflectance across changes in illumination, as well as across changes in other contextual variables (e.g., object shape, position and pose). Our study shows that population representations early in visual cortex (V1 and V4) combine information about the disk luminance and context so that a subsequent high-precision linear readout could lead to a representation of lightness consistent with the paint-shadow effect. Our work leaves open the question of whether such a readout is in fact deployed by the visual system, as there are also high-precision readouts that are inconsistent with perceived lightness. In addition, it is an open question as to whether a single fixed readout can accommodate lightness constancy with respect to contextual changes beyond the paint-shadow manipulation we studied.

Our stimuli had the property that they equate the luminance of the local and global surround of the disks. They thus seem likely to silence a number of retinal mechanisms that contribute to lightness perception more generally: contrast coding and light adaptation. We designed the stimuli to emphasize the role of cortical processing in our measurements, and our work does not characterize the contributions of contrast coding and light adaptation, nor distinguish luminance coding from contrast coding. It would be interesting in future work to study more general of stimulus manipulations with our methods.

### Is it possible to positively identify the neuronal mechanism for lightness perception given experimentally feasible data sets?

Our results speak to how the representations of luminance in neuronal populations in V1 and V4 meet the three criteria we described for a candidate neuronal mechanism underlying the checker-shadow illusion. First, we showed that both areas encode luminance (or equivalently for our stimuli, contrast) with relatively high fidelity, and that the neural sensitivity of both V1 and V4 populations depends on disk luminance in a manner qualitatively similar to the dependence revealed by human psychophysics.

Second, we showed that the representations of luminance and context interact in the neuronal population.

With respect to the third criterion, that plausible methods of decoding neuronal lightness from population responses should result in the luminance of shadow stimuli being read out as higher in lightness than that of corresponding paint stimuli, our results show that for many sessions in both V1 and V4, it is possible to construct high precision luminance decoders that result in a paint-shadow effect that is similar to our psychophysical results (e.g. all decodings from V1 or V4 whose range bars cross the solid black line in Figure 8B, C). It would be tempting to declare “victory” at this juncture and conclude that the existence of such decoders implies that population responses these areas form the neural basis of the paint-shadow effect. However, we also found a substantial number of high precision luminance decoders that are accompanied by the opposite paint-shadow effect, and even more that are consistent with no paint-shadow effect at all. Thus, conclusions about the relation between the neural population responses in V1 and V4 are contingent on assumptions about how the information carried by the population is read out. Our current data do not test these assumptions. We emphasize that this uncertainty was not a foregone conclusion: for some sessions, we do find responses where the range of decoder paint-shadow effects is sufficiently narrow to allow strong statements about the population recorded in that session. Had all sessions revealed decoders that were consistent in this manner, our data would have supported stronger conclusions.

Given the above, an important lesson from this study is that the neuronal weightings corresponding to high-precision lightness decoders (or likely decoders of any visual feature from experimentally feasible numbers of units) are far from unique. That is, the weighting can change substantially across decoders that have close to equal precision, resulting in a large range of (e.g.) paint-shadow effects.

Why were we unable to conclusively identify a neuronal mechanism for the paint-shadow effect? There are at least four possibilities:

1. Monkeys do not experience the illusion. This seems unlikely given the similarity of monkey and human vision, but training monkeys to do a lightness task using the current stimuli would be required to positively rule out this possibility (see Huang et al., 2002).
2. We were looking in the wrong brain areas. Previous work suggests that early visual areas are involved with lightness perception (Rossi and Paradiso, 1996; Rossi et al., 1996; Rossi and Paradiso, 1999; Kinoshita and Komatsu, 2001; MacEvoy and Paradiso, 2001; Haynes et al., 2004; Perna et al., 2005; Roe et al., 2005; Cornelissen et al., 2006; Boyaci et al., 2007; Pereverzeva and Murray, 2008; Boyaci et al., 2010), but the paint-shadow effect could in principle be revealed more clearly if we had applied our methods to characterize population activity in another, potentially downstream area (such as inferotemporal cortex; for example, see Leopold and Logothetis, 1996; Sheinberg and Logothetis, 1997). Indeed, our results suggest that the paint-shadow effect is set up by the representation of lightness in early visual cortex but finalized by the particular way that other parts of the brain process and read out this information.
3. We focused on the wrong subsets of neurons. Previous single unit studies of the neural basis of lightness optimized stimulus features (e.g. size and location) to match the tuning of the unit under study. Our multi-neuron recording approach made that impossible, and as a result, the majority of the units we recorded had tuning features that were suboptimal for the particular visual stimuli we presented. To explore this possibility, we made pseudo-populations consisting only of the units in which the absolute value of their luminance indices (computed as in Figure 4) were in the top decile. Unsurprisingly, these pseudo-populations encoded more luminance information than the actual recorded populations. As with the recorded populations, however, high quality lightness decodings were accompanied by a wide range of paint-shadow effects. This result is consistent with the observation that units with high magnitude luminance indices had paint-shadow indices of both signs (Figure 4C). That is, our recordings contained units that respond more to high luminance and either more to paint than shadow stimuli, or vice versa. Therefore, even if we recorded only from subpopulations of units for which the stimuli were optimized, we likely would have reached the same conclusions. It is also possible that our electrode arrays missed critical sub-populations of neurons, possibly because the most relevant neurons are located in cortical layers not adequately sampled by our electrodes. 4) V1 and V4 contain neuronal mechanisms that are the direct correlate of the paint-shadow effect, but we recorded from too few neurons to reveal these mechanisms, or applied the wrong analysis methods. Indeed, the lightness decoding weights we obtained are likely different than the ones the monkey uses. There are many possible reasons for this: we recorded from a very small subset of the neurons that respond to these stimuli; the luminance sensitivity, context dependence, and trial-to-trial variability of many neurons are correlated (meaning that even the truly optimal weights are non-unique); the monkey may use a non-linear decoder whose behavior differs in fundamental ways from that of the linear decoder we considered; the neuronal responses might be modulated by attention or motivation if our monkeys had been performing a luminance discrimination task; etc. Recording from all of the relevant neurons is not feasible; there are likely many thousands of neurons in multiple areas that respond to these stimuli, and constructing decoders with that many neurons would require a number of trials that is several orders of magnitude larger than is experimentally feasible.
4. Whether physiological data of the sort we recorded can ever positively identify the neuronal mechanisms underlying phenomena such as the paint-shadow effect remains an interesting question for future work. Consistent with previous work (Rossi and Paradiso, 1996; Rossi et al., 1996; Rossi and Paradiso, 1999; Kinoshita and Komatsu, 2001; MacEvoy and Paradiso, 2001; Haynes et al., 2004; Perna et al., 2005; Roe et al., 2005; Cornelissen et al., 2006; Boyaci et al., 2007; Pereverzeva and Murray, 2008; Boyaci et al., 2010), we did observe a small number of units in both V1and V4 that exhibited strong sensitivity to both luminance and context (see Figure 4). It remains possible that this subset of neurons plays a special role in lightness perception, but our data are equally consistent with the possibility that they do not. One approach to address this possibility, as well as the issue raised in 1) above, would be to train the monkeys to do a lightness discrimination task. We could then try to infer decoders that predict behavior and examine the units whose firing strongly influenced the decoders. Successful identification of this type of choice-based decoder might require a richer behavior than the typical two alternative forced choice discrimination task. In addition, the power of this method might be improved by increasing the number of stimulus dimensions varied in the experiments.

### Neuronal population measures can reveal representations of sensory or cognitive factors for which individual neuronal responses are heterogeneous

Many studies have found that the responses of single neurons in early visual cortex are context dependent (Rossi and Paradiso, 1996; Rossi et al., 1996; MacEvoy et al., 1998; Rossi and Paradiso, 1999; Kinoshita and Komatsu, 2001; MacEvoy and Paradiso, 2001; Friedman et al., 2003; Roe et al., 2005), but linking neuronal activity to lightness perception has proven difficult. Our approach differs from previous investigations in that we recorded from populations of neurons using a stimulus that was not optimized for the specific neurons under study. This experimental situation is more analogous to natural vision, where different aspects of a complex stimulus fall on the receptive fields of different neurons. Using these stimuli, our results demonstrate that neuronal responses to luminance and context are extremely heterogeneous (Figure 4; see also Bushnell et al., 2011; Sani et al., 2013), even as early as primary visual cortex. Furthermore, in response to these stimuli, many individual units were not very sensitive to luminance (e.g. many luminance indices were close to 0; Figure 4). None-the-less, the best luminance decoders drew non-trivially on the responses of many units. These results suggest that lightness, and likely many other perceptual phenomena, may arise from the readout of activity across large neuronal populations. Understanding the nature of these population representations in sensory cortices and how they are read-out by downstream areas will require recording from, and analyzing the responses of, neuronal populations as a whole in response to multiple dimensions of stimulus variation, likely coupled with behavioral measures (see also Shapley and Hawken, 2011).

Together, our results suggest that information about luminance and context is encoded in large neuronal populations in V1 and V4 in a manner that could, but does not necessarily, account for the paint-shadow effect.

## Footnotes

1 The null value varies slightly from session to session, depending on how many stimuli of each disk luminance were presented in that session. The reported value of 0.24 is the mean over sessions where the decoded RMSE was better than 0.20.

2 In preliminary analyses, we also explored fitting the data with an intercept parameter and the line constrained to have a slope of 1. These fits were of about the same quality as the slope only fits, and we chose the slope only fits for the theoretical reason that these describe the gain-change computation required to achieve lightness constancy across a change of illuminant intensity.

3 This fitting choice parallels the way we obtained the psychophysical paint-shadow effect. As with the psychophysical data, we have not pursued a detailed comparison of different functional forms for fitting the relationship illustrated by Figures 7B and D.

## Acknowledgments

We thank Joshua Alberts for assistance with recordings, Karen McCracken for technical assistance, and Amy Ni for helpful comments on an earlier version of the manuscript. The authors are supported by NIH grants 4R00EY020844 – 03 and R01 EY022930 (MRC), R01 EY10016 (DHB), a training grant slot on NIH 5T32NS7391-14 (DAR), a Whitehall Foundation Grant (MRC), a Klingenstein-Simons Fellowship (MRC), two grants from the Simons Foundation Collaboration on the Global Brain (MRC, 709862; DHB, 324759), a Sloan Research Fellowship (MRC), and a McKnight Scholar Award (MRC). This work was also supported by core grants from the NIH, P30 EY008098 (MRC) and P30 EY001583 (DHB).

## References

Adelson EH (2000) Lightness Perception and Lightness Illusions. In: The New Cognitive Neurosciences, 2 Edition (Gazzaniga M, ed), pp 339–351. Cambridge, MA: Press, MIT.

Boyaci H, Fang F, Murray SO, Kersten D (2007) Responses to Lightness Variations in Early Human Visual Cortex. Current Biology 17:989–993.

Boyaci H, Fang F, Murray SO, Kersten D (2010) Perceptual grouping-dependent lightness processing in human early visual cortex. Journal of vision 10:4.

Brainard DH (1997) The Psychophysics Toolbox. Spatial vision 10:433–436.

Brainard DH, Pelli D, Robson T (2002) Display characterization. In: Encylopedia of Imaging Science and Technology (Hornak J, ed), pp 172–188. New York: Wiley.

Bushnell BN, Harding PJ, Kosai Y, Bair W, Pasupathy A (2011) Equiluminance cells in visual cortical area v4. J Neurosci 31:12398–12412.

Cornelissen FW, Wade AR, Vladusich T, Dougherty RF, Wandell BA (2006) No Functional Magnetic Resonance Imaging Evidence for Brightness and Color Filling-In In Early Human Visual Cortex. Journal of Neuroscience 26:3634–3641.

Corney D, Haynes JD, Rees G, Lotto RB (2009) The brightness of colour. PLoS ONE 4:e5091.

Cowley BR, Kaufman MT, Butler ZS, Churchland MM, Ryu SI, Shenoy KV, Yu BM (2013) DataHigh: graphical user interface for visualizing and interacting with high-dimensional neural activity. J Neural Eng 10:066012.

DeAngelis GC, Uka T (2003) Coding of horizontal disparity and velocity by MT neurons in the alert macaque. Journal of neurophysiology 89:1094–1111.

Friedman HS, Zhou H, von der Heydt R (2003) The coding of uniform colour figures in monkey visual cortex. The Journal of physiology 548:593–613.

Fusi S, Miller EK, Rigotti M (2016) Why neurons mix High dimensionality for higher cognition. Current Opinion in Neurobiology 37:66–74.

Gilchrist A (2006) Seeing Black and White. Oxford: Oxford University Press.

Gold JI, Shadlen MN (2007) The neural basis of decision making. Annual review of neuroscience 30:535–574.

Haynes JD, Lotto RB, Rees G (2004) Responses of human visual cortex to uniform surfaces. Proceedings of the National Academy of Sciences of the United States of America 101:4286–4291.

Heekeren HR, Marrett S, Ungerleider LG (2008) The neural systems that mediate human perceptual decision making. Nat Rev Neurosci 9:467–479.

Hillis JM, Brainard DH (2007) Distinct mechanisms mediate visual detection and identification. Curr Biol 17:1714–1719.

Huang X, Paradiso MA (2008) V1 response timing and surface filling-in. Journal of Neurophysiology 100:539–547.

Huang X, MacEvoy SP, Paradiso MA (2002) Perception of brightness and brightness illusions in the macaque monkey. The Journal of neuroscience 22:9618–9625.

Hung CP, Ramsden BM, Roe AW (2007) A functional circuitry for edge-induced brightness perception. Nature neuroscience 10:1185–1190.

Kingdom F, Prins N (2010) Psychophysics: A Practical Introduction.

Kingdom FA (2011) Lightness, brightness and transparency: A quarter century of new ideas, captivating demonstrations and unrelenting controversy. Vision Research 51:652–673.

Kinoshita M, Komatsu H (2001) Neural representation of the luminance and brightness of a uniform surface in the macaque primary visual cortex. Journal of neurophysiology 86:2559–2570.

Leopold DA, Logothetis NK (1996) Activity changes in early visual cortex reflect monkeys’ percepts during binocular rivalry. In: Nature, pp 549–553.

MacEvoy SP, Paradiso Ma (2001) Lightness constancy in primary visual cortex. Proceedings of the National Academy of Sciences of the United States of America 98:8827–8831.

MacEvoy SP, Kim W, Paradiso Ma (1998) Integration of surface information in primary visual cortex. Nature neuroscience 1:616–620.

Majaj NJ, Hong H, Solomon EA, DiCarlo JJ (2015) Simple Learned Weighted Sums of Inferior Temporal Neuronal Firing Rates Accurately Predict Human Core Object Recognition Performance. J Neurosci 35:13402–13418.

Pagan M, Urban LS, Wohl MP, Rust NC (2013) Signals in inferotemporal and perirhinal cortex suggest an untangling of visual target information. Nat Neurosci 16:1132–1139.

Paradiso MA, Blau S, Huang X, MacEvoy SP, Rossi AF, Shalev G (2006) Lightness, filling-in, and the fundamental role of context in visual perception. Progress in Brain Research 155:109–123.

Pelli D (1997) The VideoToolbox software for visual psychophysics: Transforming numbers into movies. Spatial vision 10:437–442.

Pereverzeva M, Murray SO (2008) Neural activity in human V1 correlates with dynamic lightness induction. Journal of vision 8:8.1–10.

Perna A, Tosetti M, Montanaro D, Morrone MC (2005) Neuronal mechanisms for illusory brightness perception in humans. Neuron 47:645–651.

Radonjić A, Brainard DH (2016) The nature of instructional effects in color constancy. Journal of Experimental Psychology: Human Perception and Performance 42:847–865.

Rigotti M, Barak O, Warden MR, Wang XJ, Daw ND, Miller EK, Fusi S (2013) The importance of mixed selectivity in complex cognitive tasks. Nature 497:585–590.

Roe AW, Lu HD, Hung CP (2005) Cortical processing of a brightness illusion. Proceedings of the National Academy of Sciences of the United States of America 102:3869–3874.

Rossi AF, Paradiso MA (1996) Temporal limits of brightness induction and mechanisms of brightness perception. Vision Research 36:1391–1398.

Rossi AF, Paradiso MA (1999) Neural correlates of perceived brightness in the retina, lateral geniculate nucleus, and striate cortex. The Journal of neuroscience 19:6145–6156.

Rossi AF, Rittenhouse CD, Paradiso MA (1996) The Representation of Brightness in Primary Visual Cortex. Science 273:1104–1107.

Sani I, Santandrea E, Golzar A, Morrone MC, Chelazzi L (2013) Selective tuning for contrast in macaque area V4. J Neurosci 33:18583–18596.

Shapley R, Hawken MJ (2011) Color in the cortex: single- and double-opponent cells. Vision Res 51:701–717.

Sheinberg DL, Logothetis NK (1997) The role of temporal cortical areas in perceptual organization. Proceedings of the National Academy of Sciences of the United States of America 94:3408–3413.

Smolyanskaya A, Ruff DA, Born R (2013) Joint tuning for direction of motion and binocular disparity in macaque MT is largely separable. Journal of Neurophysiology 110:2806–2816.

Vladusich T, Lucassen MP, Cornelissen FW (2006) Do cortical neurons process luminance or contrast to encode surface properties? Journal of neurophysiology 95:2638–2649.

Yu BM, Cunningham JP, Santhanam G, Ryu SI, Shenoy KV, Sahani M (2009) Gaussian-process factor analysis for low-dimensional single-trial analysis of neural population activity. J Neurophysiol 102:614–635.

